# Hypothesis-driven interpretable neural network for interactions between genes

**DOI:** 10.1101/2024.04.09.588719

**Authors:** Shuhui Wang, Alexandre Allauzen, Philippe Nghe, Vaitea Opuu

## Abstract

Mechanistic models of genetic interactions are rarely feasible due to a lack of information and computational challenges. Alternatively, machine learning (ML) approaches may predict gene interactions if provided with enough data but they lack interpretability. Here, we propose an ML approach for interpretable genotype-to-fitness mapping, the Direct-Latent Interpretable Model (D-LIM). The neural network is built on a strong hypothesis: mutations in different genes cause independent effects in phenotypes, which then interact via non-linear relationships to determine fitness. D-LIM predicts interpretable genotype-to-fitness maps with state-of-the-art accuracy for gene-to-gene and gene-to-environment perturbations in deep mutational scanning of a metabolic pathway, a protein-protein interaction system, and yeast mutants for environmental adaptation. The hypothesis-driven structure of D-LIM offers interpretable features reminiscent of mechanistic models: the inference of phenotypes, identification of trade-offs, and fitness extrapolation outside of the data domain.

## 1 Background

Genomic mutations induce changes in fitness, or the degree of adaptation, of one organism to its environment. Mutations first affect certain parameters, *e*.*g*. the stability of specific proteins. Then, these parameters couple via metabolic or regulatory networks, ultimately determining the fitness of organisms. Typically, studies of genotype-to-fitness maps are done using large-scale experimental screens called deep mutational scans [1, 2], which consist of mutating genes and measuring the resulting fitness. With the progress of DNA-editing technologies combined with next-generation sequencing, these experiments can yield millions of fitness measurements [3]. In addition to being critically important for understanding molecular adaptations, genotype-to-fitness maps are central to biomedical research in genetic diseases, the spread of infectious diseases, and drug resistance [4, 5].

Mathematical models of genotype-to-fitness maps are of the form *F*(*g*_*i*_), where the fitness *F* is a function of the mutations *g*_*i*_ [6]. Mechanistic models typically consider two distinct levels *g* and *P* so that *F* = *f* (*P*(*g*_*i*_)). Indeed, mutations in a gene are first associated with parameters or phenotypes *P*, for example, the affinity of an enzyme for its substrate. These parameters are determined by how biomolecules fold, bind, and other biophysical properties. While some of these can be predicted with ML since recently [7], effects of mutations on properties such as catalysis cannot yet be predicted reliably. Thus, in practice, *P* is inferred by fitting experimental measurements. Then *f* is chosen among a family of parametric models, based on our knowledge of detailed biochemical processes, namely metabolic networks, regulatory networks, or any other physical interaction that is already known [8].

For instance, in the study by Kemble *et al* [9], the authors performed a mutational scan of 2 genes interacting via a metabolic pathway. They derived a mechanistic model *f*(*A, B*) based on Michaelis-Menten kinetics and the structure of the pathway. *A* and *B* are the enzymatic activities, which together determine fitness via *f*, where *f* itself depends on other parameters that account for cell physiology properties (gene expression cost, metabolic flux, toxicity). Because the choice of model family dictates the role of each parameter, and these parameters are directly associated with a gene, this model is interpretable. However, to find the parameters as a function of mutations, the authors used 10^8^ Monte Carlo optimization steps to find the most consistent mapping, under the constraint of the family *f*. While being highly interpretable, finding the correct set of hypotheses—which parametric family to use—and formalizing them into a mathematical model is challenging. Furthermore, there is no evident framework to optimize their parameters. Thus, this approach cannot be easily generalized to other similar experiments, especially in the context of larger or lesser characterized gene networks.

In contrast, machine learning (ML) does not necessitate biological hypotheses to construct models. Modern approaches, such as neural networks (NNs), capitalize on the increasing volume of data, resulting in a significant enhancement in prediction accuracy, particularly in the context of experimental mutational scanning of proteins [1]. Indeed, NNs are universal approximators [10], which means that, instead of manually selecting the parametric family, the learning process autonomously determines the choice of *f*(*h*), where *h* is a multi-dimensional latent vector representation of the genotype. Furthermore, parameter optimization is highly efficient via the backpropagation algorithm. However, interpreting how the representation influences the predictions and its relationship with phenotypes is unclear. In ML, linear regression models are considered to be the most interpretable, but they offer limited predictive power [11], especially in the context of interactions between mutations, which are non-linear.

A recent trend has focused on making machine learning (ML) models more interpretable while maintaining their predictive power. In the context of deep mutational scan experiments, the LANTERN approach constrains the latent representation of mutations, represented by a vector *h*_*i*_, to be additive, thereby facilitating the interpretation of the model parameters, where two representation vectors pointing in the same direction should have similar effects, while the magnitude of these vectors informs on the strength of the mutation [12]. Nevertheless, the specific role of the dimensions composing the coordinates *h*_*i*_ remains unclear regarding predictions or phenotypes. From a biophysical perspective, Tareen *et al*. introduced MAVE-NN, which simplifies the development of mechanistic models for genotype-to-phenotype maps. In MAVE-NN [13], users specify the type of map—ranging from additive to NN-based—between the genotype and a latent phenotype. Subsequently, the inferred latent phenotype is transformed into fitness through a stochastic process. As a result, while the latent phenotype is never directly measured, it can be interpreted as an effective phenotype. This approach presupposes that a single phenotype is influenced by all genotypes. Faure and Lehner further developed this line of reasoning by employing multiple latent phenotypes [14]. In a different approach, GenNet introduced an interpretable NN designed to study systems of genes or metabolic pathways [15]. Its architecture is informed by *a priori* biological knowledge, with neuron connectivity defined by gene annotations, pathways, cellular expression, and tissue expression. However, this method necessitates rich information about the system under study.

Here, we present a hypothesis-driven model for gene-gene interactions, which learns from genotype-to-fitness measurements and infers a genotype-to-phenotype and a phenotype-to-fitness map. The model uses a neural network *f* built upon the hypothesis *g*_*i*_ → *φ*_*i*_ that mutations *g*_*i*_ do not interact when determining the inferred phenotypes *φ*_*i*_, but this phenotype interact via a non-linear phenotype-to-fitness relationship *f*(*φ*_1_, *φ*_2_, …). Thus, our model encodes a strong hypothesis on the structure of mechanistic relationships, namely which gene affects which phenotype, but without requiring any further *a priori* knowledge, such as the values of the phenotypic parameters involved in these relationships. However, our inferred phenotypic parameters *φ* may eventually be measured experimentally, allowing us to map our inference through a measurement process *ω*(*φ*) = *ϕ*, where *ϕ* is a true measured phenotype. The ability of the model to fit the data under such constraint informs us on the validity of our hypothesis while inferring the phenotype-to-fitness relationship, the phenotypic values, and the mutations corresponding to these values.

We first show that our model, due to its architecture, offers interpretable features of mechanistic models: the inference of phenotypic proxies, the identification of phenotypic trade-offs, and the extrapolation of the fitness prediction outside from the training data domain. Next, we applied D-LIM on three mutational scanning experiments datasets, where despite the stronger constraints of its architecture compared to existing models, our model reaches state-of-the-art accuracy in fitness prediction. The resulting inferred phenotype-to-fitness landscapes are both accurate and readily interpretable.

## 2 D-LIM

D-LIM is a neural network (NN) designed to learn genotype-to-fitness maps from a list of genetic mutations and their associated fitness when distinct biological entities referred to here as ‘genes’ for simplicity (Fig 1A). We describe in the next section its architecture, while in the following section, we demonstrate in simulated datasets that D-LIM enables us to recover phenotypic values. Finally, we show that our model generalizes beyond already explored mutations, akin to mechanistic models.

**Figure 1:**
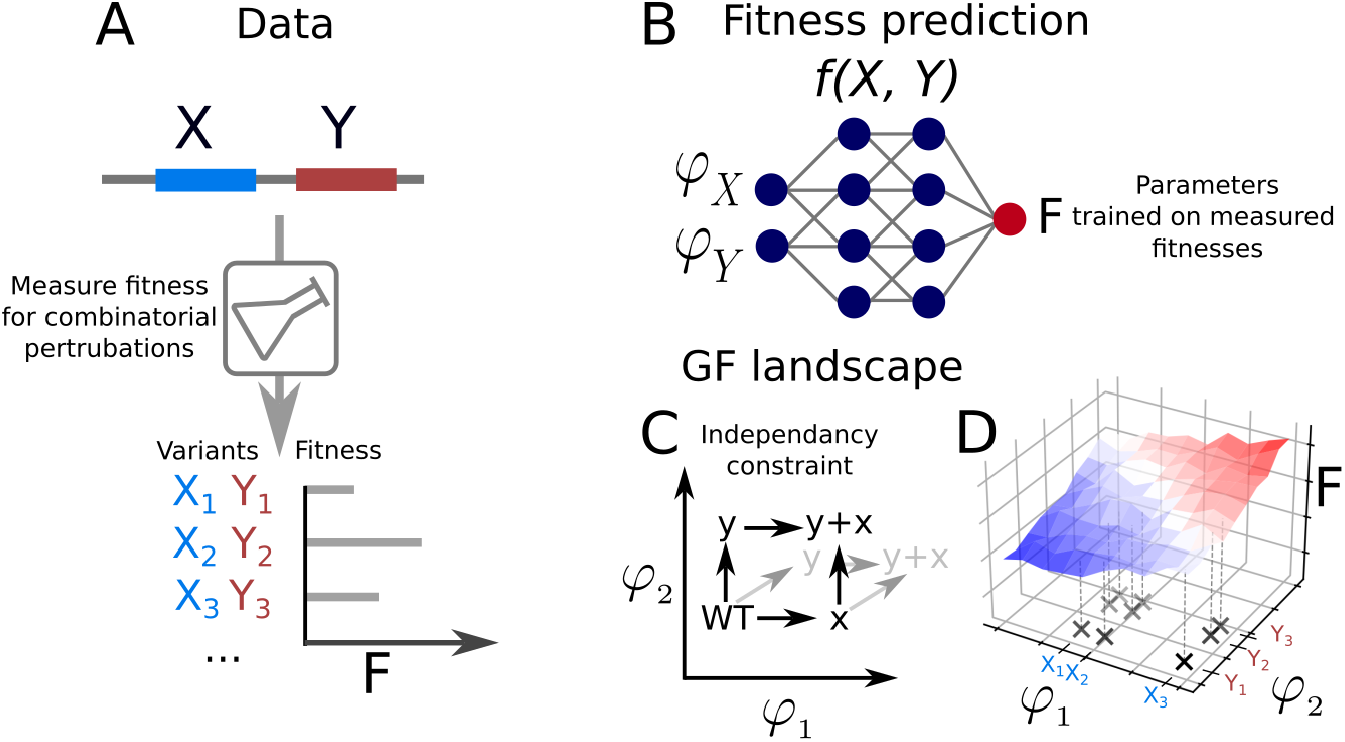
Overview of the D-LIM framework. A) Data acquisition example for gene pair X and Y with fitness measurements across mutation combinations. B) D-LIM architecture. D-LIM takes one value *φ* per gene variant and processes it through a feed-forward neural network to predict the fitness of these two gene variants. C) Inferred phenotype space. Each dimension represents a gene, with gene variant coordinates *φ* obtained during D-LIM training. Consequently, mutations in such a constrained space are independent (represented by the dark arrows). If mutations did not affect phenotypes independently, they would vary along all dimensions at once (as represented by the grey arrows). D) The constrained phenotype space. The fitness is mapped to the constrained phenotype space where combinations of gene variants are organized in a grid.

### 2.1 D-LIM architecture

The model comprises three levels of description: mutations (genotypes), phenotypes, and fitness. Phenotypes are understood here in a very general sense as fitness-determining parameters, which may represent low-level molecular properties (e.g. binding), and higher-level measurable properties (e.g. metabolic flux). The space spanned by the phenotypic values is termed the phenotype space. We connect these levels of description via a genotype-to-phenotype map and a phenotype-to-fitness map. The genotype-to-phenotype map encodes the hypothesis that mutations participating in different genes do not interact when determining phenotypes, though mutations within the same gene can interact. In the NN, this corresponds to associating each mutation with a single phenotypic value, while the phenotype-to-fitness map is learned using an NN (Fig 1B), leading to a relationship of the form 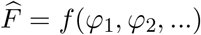.

The training of D-LIM fitting consists of two steps: (1) initializing the genotype-to-phenotype map by assigning initial values to *φ*_*i*_, and (2) simultaneously training all parameters, including the *φ*_*i*_.

The initialization starts with the assumption that mutations with the same phenotypic value yield similar fitness values in matching genetic background (in combination with the same mutations). We called this procedure spectral initialization, which for one gene follows these steps: (i) compute similarities between mutations using fitness values from the training data; (ii) extract a representation that encapsulates relationships from these similarities via spectral decomposition; (iii) initialize the values of the *φ* using this representation, see section 5.1.

Once the genotype-to-phenotype is initialized, we simultaneously train all the NN and refine the *φ*_*i*_ parameters using the Adam optimizer [16] in the Pytorch framework [17]. The parameters are updated using the Negative Gaussian Log-likelihood loss function which accounts for the measurement noise [16]. To favor a smooth landscape, we applied *L*_2_ regularization to the *φ*. During training, D-LIM refines the phenotypic values while simultaneously learning to predict fitness(Fig SI2).

Due to the gene-to-phenotype constraints, D-LIM is more rigid than existing ML approaches. For instance, in LANTERN, mutations are represented in an unconstrained latent space, where mutations may span multiple dimensions, without being linked to any specific biological entity (Fig 1C).

### 2.2 Phenotype inference

In our genotype-to-phenotype map, genes are assumed to act independently in determining the phenotypic value. While our phenotypes *φ* are not measured but inferred, they are thought to be proxies for measurable biological phenotypes. We now show on simulated data that using our hypothesis infers phenotypes related to the true underlying phenotypes.

To perform our test, we used three models to simulate datasets: (i) a mechanistic model of a two-genes metabolic pathway [9], we called here Kemble model; (ii) a tilted Gaussian used in evolutionary dynamics (called Fisher geometric model) [18]; (iii) A gene regulatory cascade with transcription factors [19]. All models consist of two genes, and for the simulation, we assigned 30 mutations per gene with phenotypic values randomly drawn in [0, 5], as shown in Fig 2. This resulted in datasets of 900 mutation pairs for which we computed fitness values with the mechanistic model, adding Gaussian noise to measurement errors. Then, for each dataset, we trained D-LIM with half of the samples. To demonstrate the importance of the spectral initialization, we trained a second model where the *φ* were randomly initialized.

**Figure 2:**
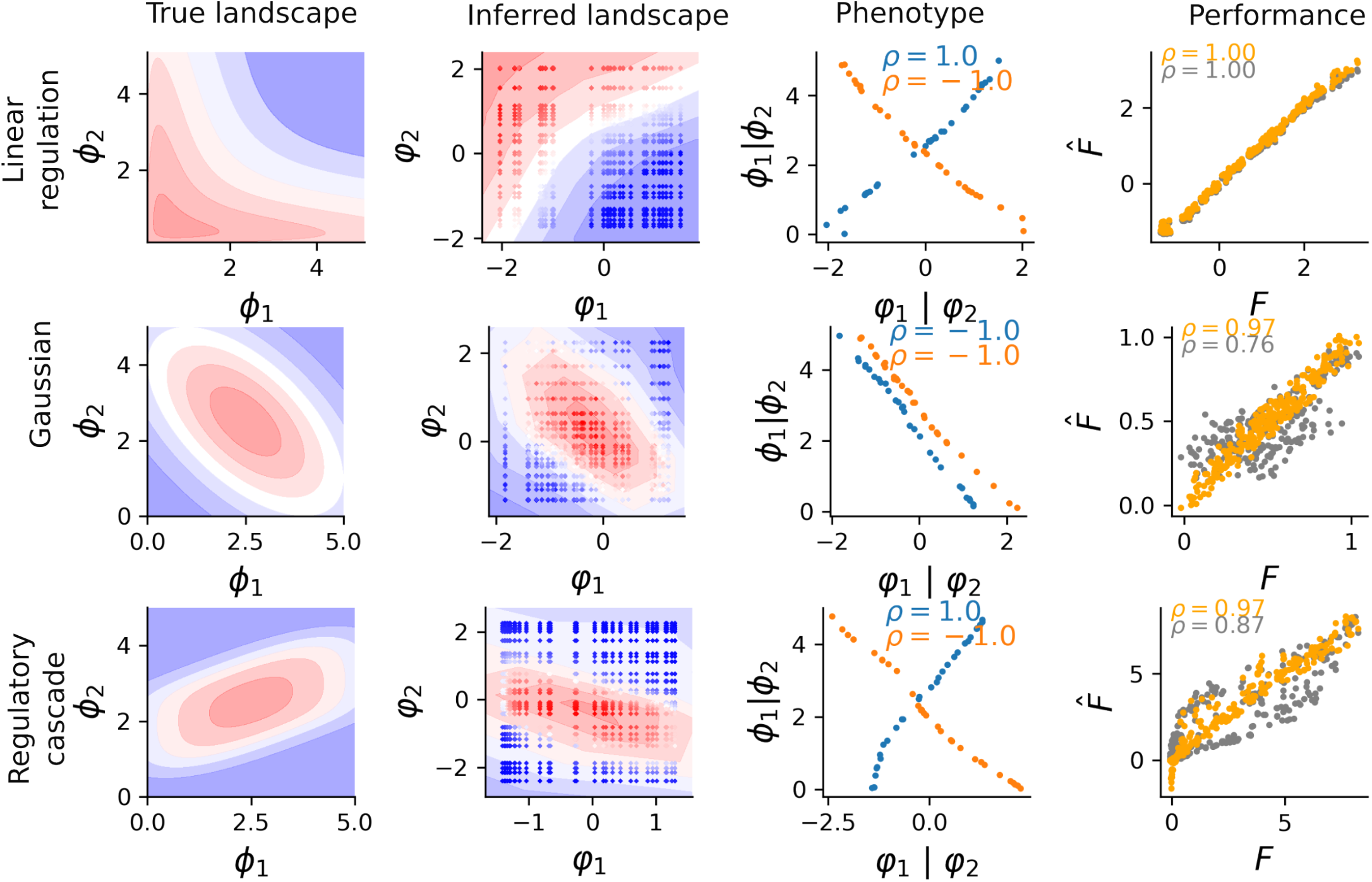
Simulated landscapes. From left to right, the plots display the theoretical fitness landscapes in a blue-to-red color gradient (first row: linear regulation of two genes, second row: geometric model with Gaussian expression, and third row: cascade model), the fitted landscape using spectral initialization; a scatter plot of true versus inferred phenotypes; a scatter plot of predicted versus true fitness values, with spectral initialization (orange) and without (grey).

For the Kemble model, we observed a good agreement with the fitness (Pearson correlation *ρ* = 0.96). While the fitness landscape spanned by the trained phenotype space is different from the original one (Fig 2), the inferred phenotypes and the corresponding measured phenotype (enzyme activity) were strongly correlated (*ρ* ≈1). We observed that both initialization strategies led to a monotonous relationship between true and inferred phenotypes and similarly high predictive power.

For the Fisher and the regulatory cascade model, we first tested the random initialization for the *φ*, which led to poor phenotypes inference with rough landscapes (Fig 2). We determined that the asymmetry of the shape is the cause of the loss of accuracy. Indeed, the shape of these models induces a strong dependency on the coordinates between the inferred phenotype axes *φ* and the fitness *F*. For example, in the non-rotated zero-centered Gaussian model *e*.*g. F* (*x, x*) = *F*(*x*, − *x*), whereas the rotated Gaussian does not allow for such symmetry. Therefore, the genotype-to-phenotype map has more optimal solutions for the non-rotated symmetrical Gaussian model (similarly for the cascade model), which may cause the loss of accuracy of D-LIM fitting.

Applying the spectral initialization showed a clear improvement in prediction accuracy and in inferred phenotypic values for the non-monotonous landscapes. With the tilted-Gaussian model, the prediction accuracy increased from *ρ* = 0.86 to *ρ* = 0.98 (see Fig 2). With the double cascade reaction, we observed consistent improvements, where the accuracy increased from *ρ* = 0.86 to *ρ* = 0.99. For the true phenotypic reproduction, we showed that the spectral initialization overcame the limitation of non-monotonous underlying landscapes *ϕ* (| *ρ*| = 1 for both models, see Fig SI4), allowing us to recover the shape of the underlying true landscape.

We have shown that our approach enables us to recover a monotonous relationship between the inferred and true phenotype, which is non-obvious given that data only provides genotype-to-fitness relationships.

### 2.3 Phenotypic measurement process for fitness extrapolation

Extrapolation of the prediction outside of the data domain is typically not expected for NN models. In contrast, mechanistic models can be fitted within a certain data domain, so that in turn, the model parameters can be used for prediction outside of the initial domain. Given that D-LIM retains some of the constraints of mechanistic approaches, we surmised that its hypothesis-driven architecture may be used for extrapolation within a certain range beyond the training data. The extrapolation strategy is based on the relationship between the inferred and actual phenotypes, requiring the measurement of some of the latter for single genes. Indeed, when experimental measurements of the phenotypes are available for some of the mutants, the inverse relationship *ω*^−1^(*ϕ*) = *φ* between inferred and actual phenotypes can be fitted by a smooth function. Fitness predictions can then be extended to new mutants by measuring their actual phenotypic values *φ* associated with single mutations, then deducing their latent phenotypic value *φ* via *ω*^−1^.

To illustrate this approach, we simulated an experiment with the Kemble model, where fitness is controlled by two phenotypes X and Y—enzyme activities by AraA and AraB. We restricted the training data to pairs of mutations yielding low fitness (black square in Fig 3A) whereas mutation combinations outside of this domain were left for validation. For the sake of comparison, we first tested a naive extrapolation strategy, directly using the model to predict fitness for combinations involving never-observed mutations. The prediction displayed poor accuracy (*ρ* = 0.59) (Fig 3B). Then, we applied our strategy using a third-order polynomial to fit the relationship between *φ*_1_, *φ*_2_, and their respective phenotypic values *ϕ*_1_, *ϕ*_2_, as shown in Fig 3C. Using the resulting phenotypes as an input for the NN yielded high fitness prediction accuracy (MSE=0.19), as shown in Fig 3B. This extrapolation technique also worked well for non-monotonous fitness functions (see Fig SI3).

**Figure 3:**
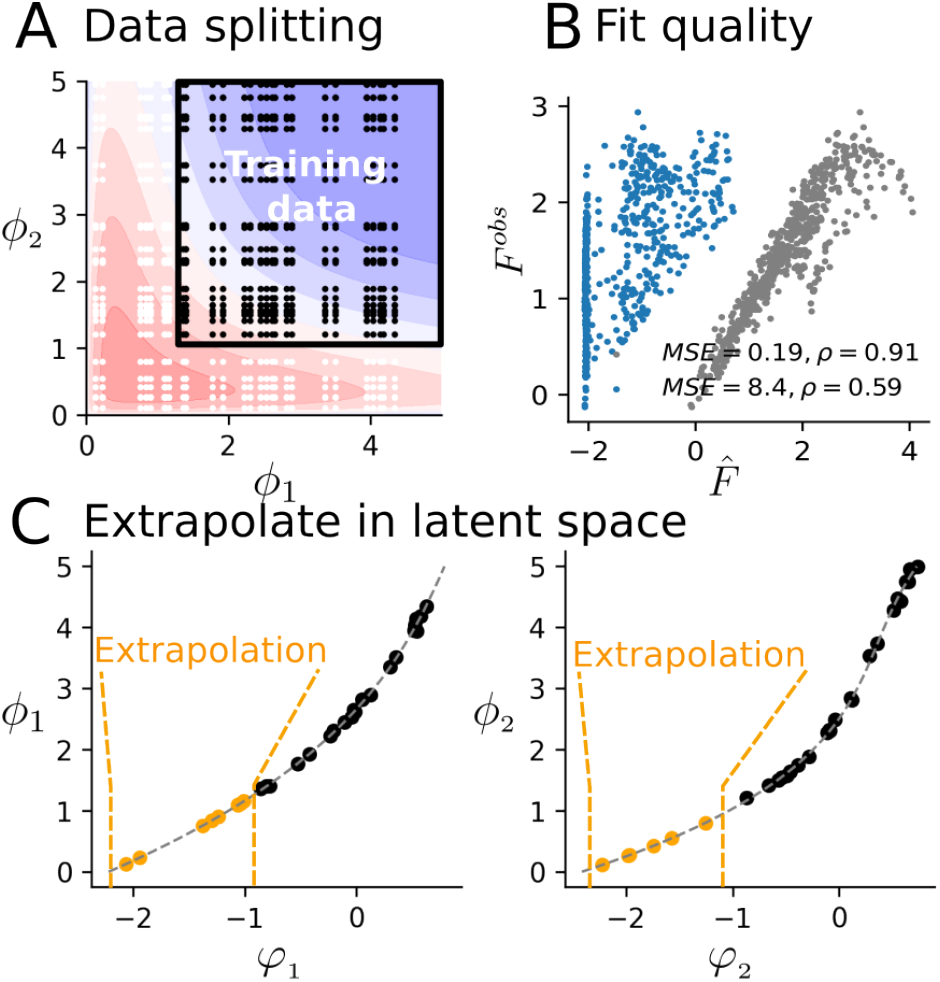
Extrapolation with un-seen mutations and their measured phenotypes. A) Fitness landscape from the mechanistic model of [9]. Black points indicate training data and white points indicate validation data. B) Fit quality. Conversion of true phenotypic values into inferred phenotypes by D-LIM for extrapolation. Blue points represent cases without additional true phenotype values, while grey points use them. C) Extrapolation using phenotypic measurements: *ϕ*_1_ (left) and *ϕ*_2_ (right). Black points (*ϕ*_1_, *φ*_1_) are used to parameterize the polynomial fit in a dotted grey line. Orange points are the converted *ϕ*_1_ and *ϕ*_2_ only seen in the validation, which are converted into inferred phenotypes using the polynomial fit.

Overall, the independence hypothesis allows D-LIM to extrapolate fitness predictions beyond the training data using phenotypic measurements, which would not have been straightforward to implement with other ML approaches. Although this extrapolation strategy requires additional measurements, it could be of practical interest because it relies on single mutation data instead of combinations of mutations.

## 3 Predictive and interpretable constrained architecture

D-LIM offers features akin to mechanistic models, going beyond typical NN approaches. We now apply D-LIM to interpret phenotypes and reveal potential trade-offs across three mutational scanning datasets: a two-gene metabolic pathway, a system of two physically interacting proteins, and yeast strains adaptation across multiple environments.

### 3.1 Revealing hidden phenotypes from gene-to-fitness

Kemble *et al*. dataset involves two genes in the metabolic pathway of L-arabinose utilization: L-arabinose isomerase (AraA) and L-ribulokinase (AraB) (See Fig 4A). These enzymes convert L-arabinose into L-ribulose-5-phosphate, which is critical for cellular growth. Mutations were systematically introduced in the promoter regions of both AraA and AraB genes, enabling comprehensive analysis of individual and combined genetic perturbations, covering a total of 111 mutations per gene promoter. The measurements were performed in three distinct environments (Env1, Env2, Env3) that differ by the concentration of inducers promoting the synthesis of both enzymes. Env1 had inducers for both genes, while Env2 and Env3 had respectively 5 ng and 200 ng of the araA inducer but without the araB inducer. The data consists of 1369 mutation combinations and the resulting fitness (measured as growth). The phenotypes are the activity of the enzymes AraA and AraB, which are not measured.

**Figure 4:**
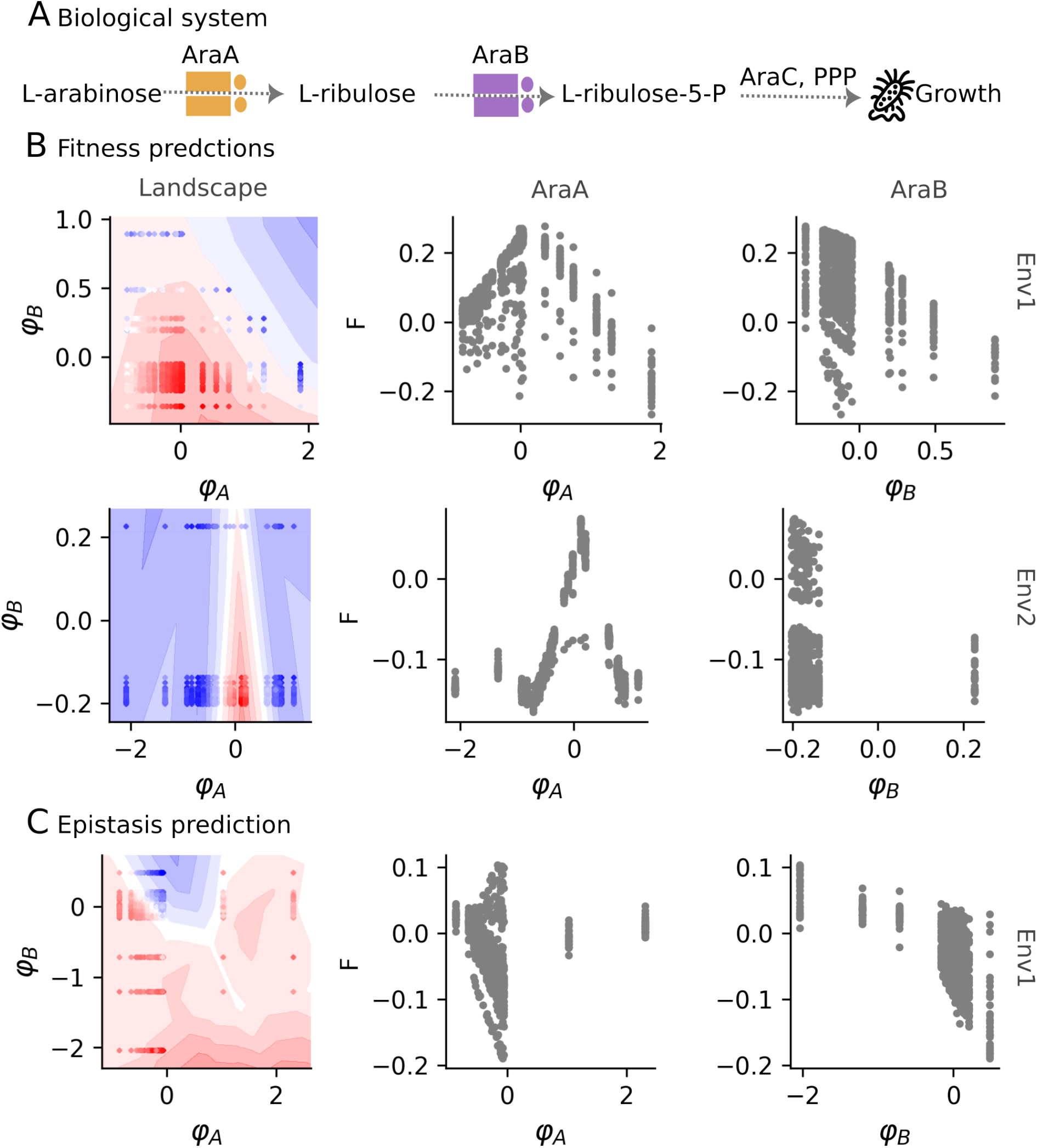
Gene-gene interaction pathway test case. A) L-Arabinose pathway of *E. coli*. B) Fitness prediction in two environments. From left to right, we show the predicted fitness landscape represented by a blue-to-red color gradient contour for the prediction, where *φ*_*A*_ and *φ*_*B*_ are the inferred phenotypes for mutations in genes AraA and AraB. The points indicate the pairs of mutations colored by the experimentally measured fitness. Next, we display the relationship between the measured fitness and the two inferred phenotypes. The first row corresponds to analyses in Env1, while the second row shows Env2. C) Epistasis prediction. This row illustrates the accuracy, landscape, and correlation between inferred phenotypes and epistasis in Env1.

We used D-LIM to predict fitness using two phenotypes *φ*_*A*_ and *φ*_*B*_, corresponding to enzymes AraA and AraB, respectively. We trained the model on 70% of the data and evaluated its accuracy on the remaining 30%—we repeated 10 times across random data splits to obtain the average correlation. In Env1 and Env2, D-LIM predictions showed strong agreement with measured fitness (*ρ* = 0.98 and *ρ* = 0.99 respectively with p-value < 10^−5^) (Fig 4B). Next, we tested the predictive accuracy on epistasis, which is the deviation in the fitness of a pair of mutations from the sum of their individual fitness *E*(*a, b*) = *F*(*a, b*) −*F*(*a*) −*F*(*b*). D-LIM predicted similarly well epistasis with *ρ* = 0.98 (p-value < 10^−5^).

In the inferred landscapes, We observed qualitative differences we quantified by analyzing the variation of fitness along the inferred phenotypes. In Env1, the fitness varies across both inferred phenotypes, indicating that both genes influence the measured fitness. However, D-LIM revealed a trade-off in AraA, that is varying *φ*_*A*_ beyond a threshold reduces the fitness (Fig 4B), whereas fitness varies monotonously along *φ*_*B*_ (|*ρ*| = 0.43 with p-value < 10^−5^). For epistasis, it varies monotonously only along *φ*_*B*_ (|*ρ*| = 0.40 with p-value < 10^−5^) (Fig 4C). In contrast, in Env2, the fitness varied along *φ*_*A*_ with a similar trade-off; but not along *φ*_*B*_ (|*ρ*| = 0.08, p-value=0.003). This means that AraB activity is anticipated to not vary between mutants. Indeed, since no AraB inducer was used in Env2, we only expected basal effects. This conclusion was reached because of the independence of phenotypes hypothesis imposed in D-LIM, which allowed us to separate the contribution of each phenotype.

Does imposing the constraint of independent phenotypes result in a trade-off with predictive accuracy? To explore this, we compared D-LIM to three analogous models that do not impose constraints on their latent spaces: a) the linear regression (LR), b) LANTERN, which uses based Gaussian process, c) the Additive Latent Model (ALM) using an NN, and d) MAVE-NN that is biophysics-based (see SI5.2 for details).

For this system, the fitness is largely additive (see Fig SI1), which can be seen from the performance of the LR (on average *ρ* = 0.97) and ALM (additive latent model, *ρ* = 0.99); so the challenge is to predict epistasis. As expected, all other models performed significantly better than LR for epistasis prediction (Fig 5). For MAVE-NN, we choose to illustrate the additive genotype-to-phenotype map offered, which essentially corresponds to a regression. For the NN-based models, we did not observe any striking loss of fitting performances when comparing D-LIM with its version with the unconstrained latent space (called ALM here).

**Figure 5:**
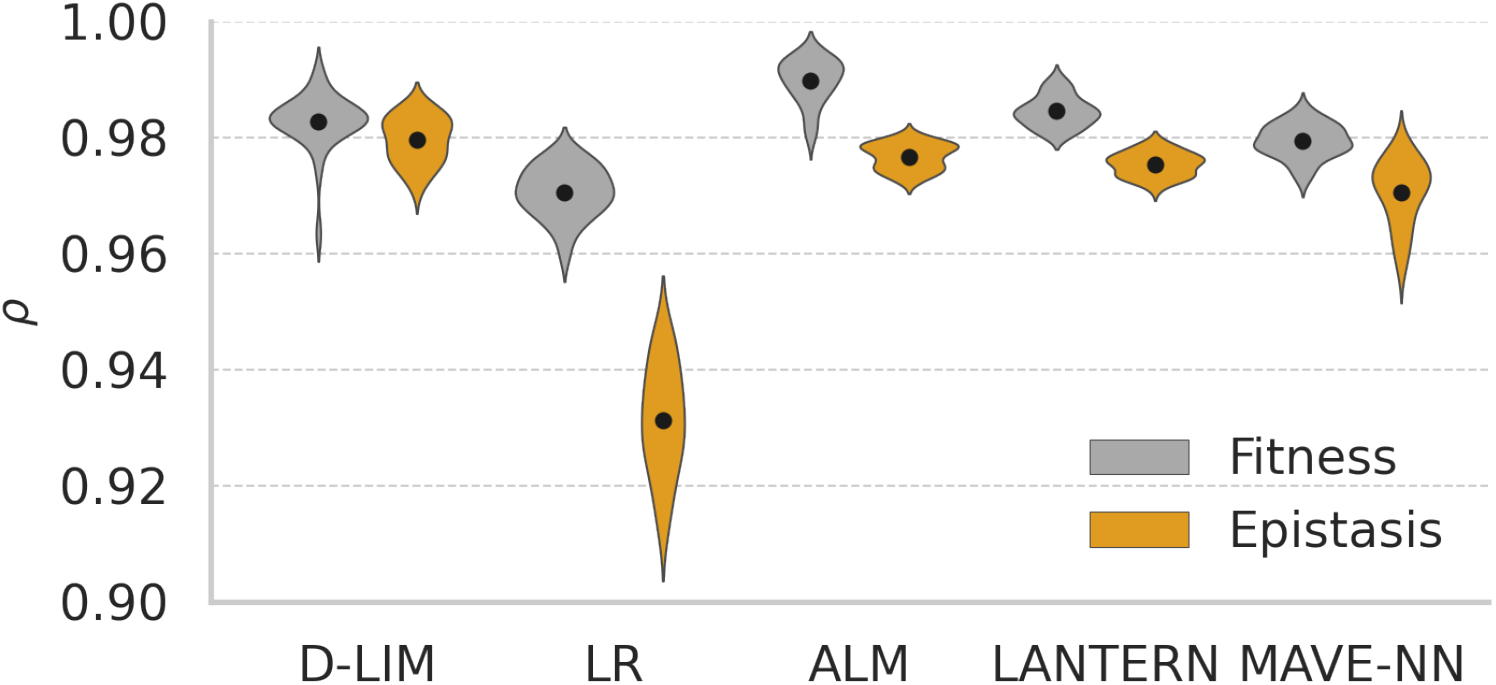
Comparing prediction across models. Violin plot of Pearson correlations for random cross-validation across 30 runs. 70% of the original data was used as training dataset and the remaining 30% as the validation set. The black dot represents the average Pearson correlation for each model on the fitness or epistasis dataset.

Finally, we showed that the inferred phenotypes are consistent with the biophysical hypotheses established earlier for this system, as implemented in the mechanistic model in [9], see details in 8.2. To validate this, we compared the inferred *φ* here with the ones inferred earlier for the mechanistic model for the Env1 since the other two are depleted with AraB inducers. Fig 6 shows that the inferred phenotypes are consistent with the ones inferred with the mechanistic model, where Spearman’s correlations *ρ*_*s*_ = 1 for both phenotypes. This demonstrates that D-LIM is capable of revealing mechanistic insights, which were explicitly modeled in the Kemble model [9].

**Figure 6:**
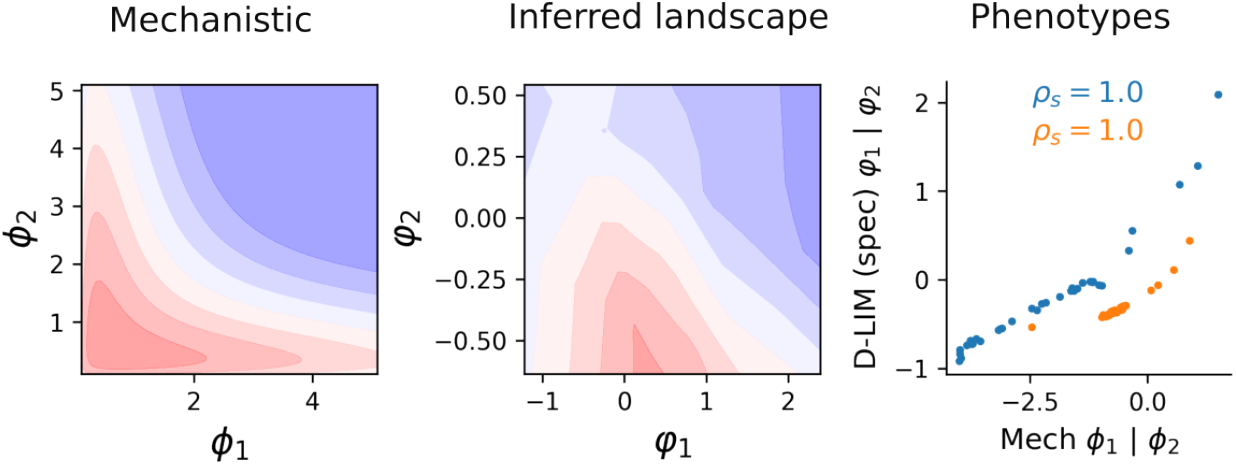
Capturing mechanistic hypotheses. (left) The mechanistic model obtained from [9] is displayed in contour. (middle) The learned landscape of D-LIM trained on all the data points of the Env1. (right) The D-LIM inferred phenotypes are compared with the phenotypes inferred earlier from the mechanistic model (extracted from [9]).

Our results showed that D-LIM performances are on par with state-of-the-art approaches for fitness and epistasis prediction. The high performances in fitness and epistasis prediction suggest that our independence of phenotype hypothesis is largely satisfied for this system, allowing D-LIM to recover mechanistic insights.

### 3.2 Interpreting mutation from physical interactions

Mutations at the genetic level influence the fitness of organisms through interactions with other genetic elements, where multiple phenotypes may be involved. In [20], Diss and Lehner quantified the effect of mutations on a system of two proteins, called FOS and JUN that interact via their leucine zipper, a 32-residue helix (Fig 7A). Consequently, the fitness measured is defined based on identified phenotypes, specifically the physical binding between the two proteins. To investigate genetic interactions, two mutations were introduced for each pair of mutants—either one mutation in each gene expressing the proteins or both mutations in one gene. The fitness for each pair of mutations depends on the binding strength of the mutated proteins quantified by the protein-protein interaction (PPI) score, resulting in over 120,000 data points. This dataset delves into two types of interactions: the first is fitness, which is mainly driven by the concentration of protein complexes, while the second is epistasis, resulting from specific structural interactions between the two protein helices.

**Figure 7:**
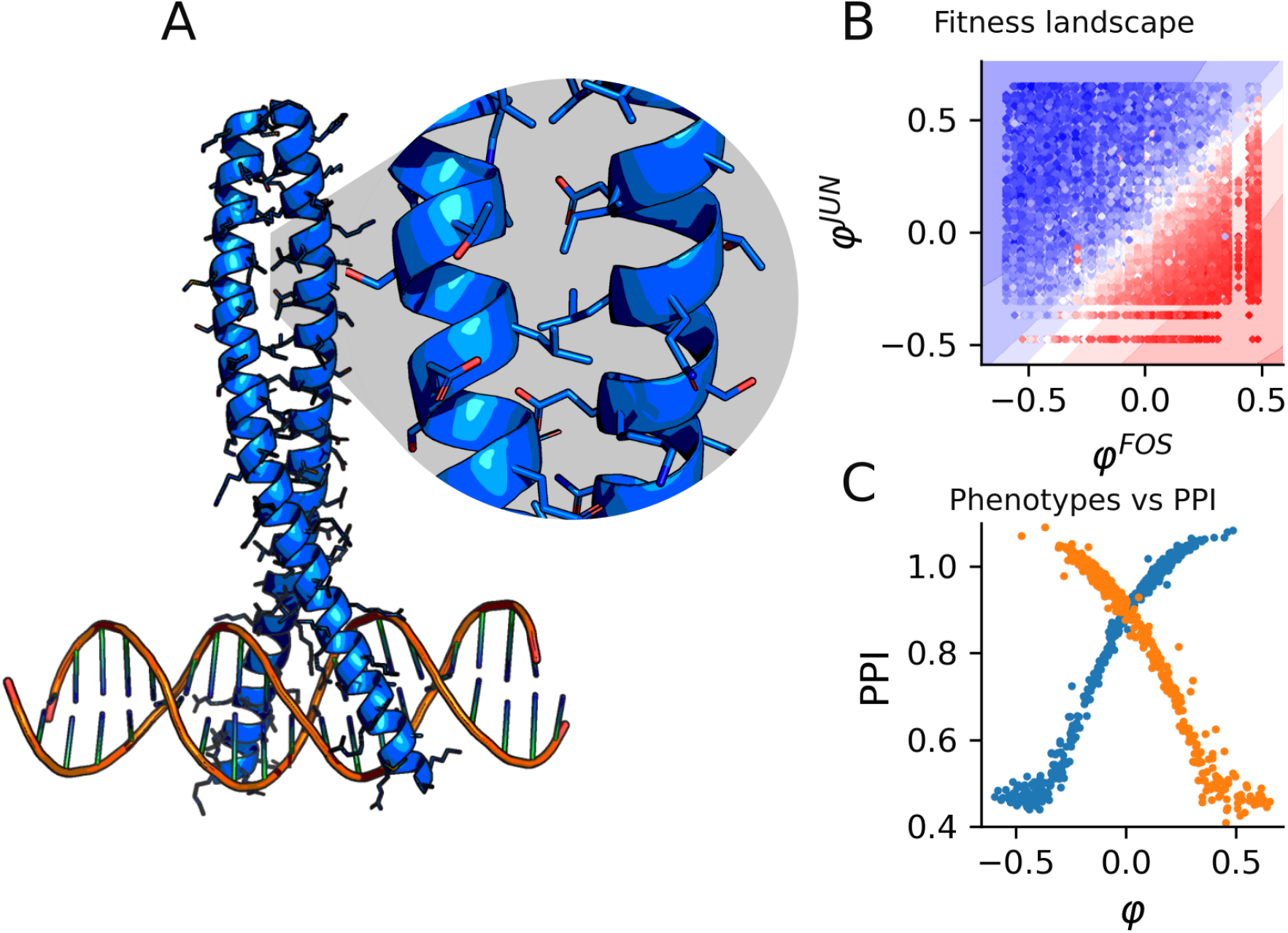
Predicting protein-protein interactions. A) JUN and FOS proteins physical interaction modulated by amino acids at the interface, with a zoom on the interacting part. B) The fitted landscape (contour) modeling the variation of fitness where dots are pairs of mutations colored by the observed fitness value whereas the contour coloring is the prediction. C) Experimental PPI scores for single mutants are represented against the inferred phenotypes from D-LIM.

We modeled this system fitness landscape using two phenotypes (*φ*^*F OS*^, *φ*^*JUN*^), one for each protein. With this dataset, the spectral initialization yielded a spectral gap *λ*_2_ *>* 0.95; therefore, we initialized the phenotypes randomly. Then, we equipped D-LIM with two layers of 64 neurons to predict the fitness *F*.

D-LIM has then been trained on 70% of the data and tested against the remaining 30%. Our prediction quality matched that of the thermodynamic model proposed alongside the data, with D-LIM explaining 93% of the variance. Fig 7B displays an additive smooth inferred landscape where fitness varies across mutations on both proteins.

We then compared the inferred *φ* values with the single mutation PPI score, obtained experimentally, which quantifies the strength of the physical interaction (Fig 7C). D-LIM consistently recovered the sigmoidal relationship between the fitness and the physical interaction, with an average *ρ* = 0.97 and *p*-value < 10^−5^. In contrast, D-LIM failed at modeling the epistasis (*ρ* = 0.23 with p-value < 10^−5^, Fig SI5), suggesting that our independence of phenotypes is not satisfied here.

While the fitness is additive and easy to predict, recovering the sigmoid relationship with the underlying protein-protein interaction is not. For the epistasis, D-LIM did not provide a good model. This result suggests that two independent phenotypes are not sufficient to describe this phenomenon and missing hidden interactions should be expected, which is consistent with earlier conclusions [20].

### 3.3 Predicting fitness variation across environments

In contrast to previous mutation-to-fitness datasets, Kinsler *et al*. [21] measured the fitness of 292 yeast strains across 45 different environments by varying factors such as the concentrations of NaCl, KCl, glucose, fluconazole, and geldanamycin in the growth medium, as well as the transfer time during the experiments.

For each environment, the fitness of all mutants was determined by monitoring their populations throughout the adaptation experiment using barcode sequencing. In total, 13,140 mutation-environment fitness measurements were obtained. Two types of environments were identified: those yielding small fitness perturbations (7,300 data points), that are referred to as subtle; and those yielding strong fitness perturbations (5,840 data points).

To apply D-LIM, we modeled the yeast mutations with one phenotype *φ*^*mut*^ and the environment with a second phenotype *φ*^*env*^. We started by analyzing the 13,140 data points of the subtle environments (organized in 80% training 20% validation). In D-LIM, we used two hidden layers of 64 neurons, and we initialized the *φ* using the spectral initialization procedure. Our results showed a strong correlation with the experiments, achieving *ρ* = 0.96 (p-value < 10^−5^). The fitness landscape inferred from the analysis confirmed the mitigated effect of the environment on fitness (Fig 8A). In contrast, D-LIM suggests the presence of a trade-off in the mutations. The mutations involved in the high-fitness region are in the IRA1 and IRA2 genes, which are partly responsible for the adaptation to glucose-limited environments [22]. For the strong perturbations, we observed a similar prediction accuracy (*ρ* = 0.92 with p-value < 10^−5^), but in contrast with above, the environments now is responsible for the largest variation in fitness (Fig 8B). Notably, the environments varied continuously across the phenotype *φ*^*mut*^, in NaCL and KCl, meaning that the inferred phenotype is sensitive to salt concentration.

**Figure 8:**
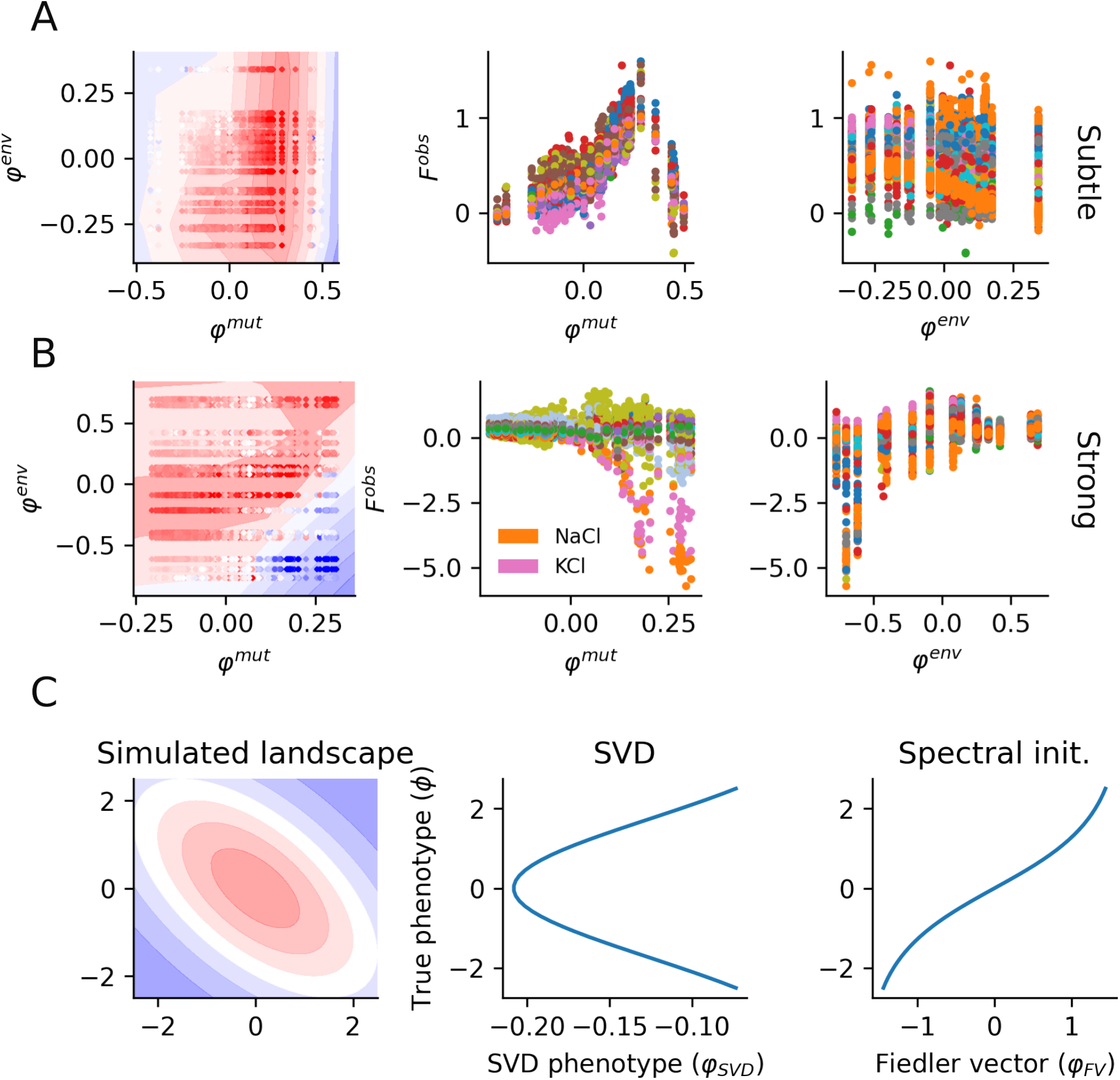
Predicting mutations across various environments. A) Subtle environments from [21]. From left to right, we show the inferred landscape, the inferred phenotype (corresponding to mutations) against observed fitness but colored by environments, and the inferred environment phenotype colored by the corresponding mutations. B) The second row presents the same plots for the Strong environment as defined in [21]. C) The SVD approach applied to the geometric Fisher model (left) is shown, where the first eigenvector (interpreted as inferred phenotypes) is plotted against the true phenotype. Finally, we compare this with the initial guess obtained from the spectral initialization procedure (Fiedler vector).

Kinsler *et al*. proposed an interpretable modeling technique based on the singular value decomposition (SVD) applied to the fitness dataset. Specifically, they constructed a matrix F, where the entries represent the measured fitness of various mutation-environment combinations. Applying the SVD to this matrix F allowed them to extract vectors representing the mutant phenotypes and vectors representing the environmental weights (respectively being the left and right eigenvectors of the SVD). Such an approach allowed them to accurately reproduce fitness using as few as eight phenotypes, corresponding to the eight most important left eigenvectors. Because 95% of fitness variability is explained by the first component, a linear relationship appears between the mean fitness and the inferred phenotypes (Fig SI6). This approach however might not represent the structure of the landscape well if this landscape is not monotonous; therefore missing potential trade-offs. To demonstrate this, we used the Fisher geometric model to create an artificial dataset from which we inferred phenotypes *φ*_*SV D*_ using the SVD, which we compared with the true phenotypes *ϕ*. Fig 8C shows that the relationship between *φ*_*SV D*_ and *ϕ* is not monotonic. In contrast, using spectral initialization, the initial guess of the *φ*_*FV*_ (the Fiedler vector) is already well related to true phenotypes, allowing us to provide a correct interpretation of the phenotypes.

To our knowledge, D-LIM is the only approach that offers a robust modeling of non-monotonous and non-symmetric fitness landscapes, enabling us to detect potential trade-offs. These trade-offs are typically missed by methods such as the SVD proposed earlier.

## 4 Discussion

Genotype-to-fitness maps are central to understanding and predicting evolutionary adaptation. These maps can be fitted with mechanistic models, which are expected to be predictive and interpretable. However, they often require prior knowledge which may be unavailable, especially in biological systems. In contrast, ML models can be predictive without knowledge of the mechanisms, but at the cost of interpretability. In this work, we proposed D-LIM, a model that encodes biological hypotheses in the ML architecture: a relationship between mutations and genes, and the constraint that one gene affects only one phenotype. D-LIM thus mimics the mathematical form of a mechanistic model, but where the phenotype-to-fitness relationship is found automatically by neural network instead of being fitted with a parametric family of biochemical functions.

D-LIM is interpretable in the classical sense of hypothesis testing, where the fitting quality assesses the hypotheses underlying the model. Specifically, using data from mutants in a metabolic pathway, we showed that D-LIM predicts epistasis between genes with the accuracy of mechanistic and pure ML models, thus validating the hypothesis that mutations in each metabolic gene mostly affect the properties of the enzyme encoded by this gene, and not other enzymes. This contrasts with mechanistic models, which may validate additional hypotheses (e.g. the functional form of enzymatic responses), and with ML models, which would predict fitness without proving or disproving any property of the metabolic pathway.

In contrast with ML, mechanistic models typically have virtues additional to pure prediction, due to their interpretability: the inference of biological parameters, the extrapolation beyond the data range, and the improvement of the integration of prior information. As D-LIM is partly constrained by the biological hypotheses used in mechanistic models, we showed that these virtues can be partly recovered.

First, D-LIM infers phenotypes that are formally the coordinates of a latent space between mutations and fitness. Although the inferred phenotypes are not *a priori* related to biological phenotypes, we demonstrated that the former could be interpreted as proxies of the latter up to a non-linear relationship. In particular, when the true phenotype-to-fitness relationship is monotonous, there is a univocal and monotonous relationship between each inferred phenotype and each biological one. For the case where the true phenotype-to-fitness is non-monotonous, we devised a learning strategy called spectral initialization: it first initializes genotype-to-phenotype maps using similarity between genotype effects, which helps the training algorithm converging toward an univocal and monotonous relationship between inferred and biological phenotypes, allowing us to detect trade-offs. These trade-offs, which are common in biological systems, are often difficult to capture. Indeed, approaches such as the SVD proposed in [21] are not capable of capturing trade-offs in the typical Fisher geometric model, whereas D-LIM, to our knowledge, is the only method tested that can do this. Other ML methods use latent spaces, where a mutation may span multiple dimensions. In LANTERN, the proximity between mutations in the latent space indicates a similarity between their effects. However, the coordinates of mutations cannot be directly interpreted as a biological property. In MAVE-NN, there is a single latent interpretable phenotype, but the approach is not suited for several phenotypes. MoCHI addresses the case of several phenotypes but does not deconvolve the effect of separate genes. In GenNet, nodes of the NN are connected according to the underlying gene network. Although this could improve interpretability, it requires extensive *a priori* biological knowledge, and the relationship with the phenotypes remained elusive.

Secondly, D-LIM yields landscapes that are consistent with mechanistic hypotheses, using only fitness measurements. With fitness measurements, we do not have access to the real phenotypic values under which our foundational hypothesis is defined. To overcome this limitation, in studies such as [9], one defines a set of hypotheses to build a mechanistic model, which links phenotypes to fitness. Using our spectral initialization, we recovered the ordering of the phenotypic values inferred earlier by their mechanistic model. Not only does this result support the mechanistic hypotheses drawn earlier, but it also shows that D-LIM retrieves such hypotheses automatically. Moreover, our procedure allows us to overcome the convergence in local minima during training, making the spectral initialization strategy applicable generally beyond the scope of this work. Importantly, the independence hypothesis underlying D-LIM provides a generic manner to use spectral initialization, where mutations in a gene are considered close when they yield similar responses when other genes vary. This inference notably shows that D-LIM can generate novel important mechanistic hypotheses, such as the existence of intermediate phenotypic values that optimize fitness.

Finally, if provided with some phenotype values, D-LIM can perform extrapolation beyond the initial training domain. Extrapolation was performed in the latent space, by fitting the relationship between the inferred versus actual phenotypes with a polynomial. Measuring the phenotype of new mutants allows one to project them into the latent space, and then compute their fitness using the NN. This may be of practical interest, as it requires measuring phenotypes for single mutations instead of combinations thereof.

Overall, our results show that introducing constraints on the form of the model enhanced the interpretability of neural networks while retaining most of their predictive power. At this stage, the hypotheses need to be determined by the user. However, it may become possible to automatically generate the mutations-to-genes mapping, ultimately allowing us to devise interpretable ML for large biological systems.

## 5 Methods

### 5.1 Spectral initialization

We now describe the application of the spectral initialization to a two-gene system, where fitness is available across pairs of mutations. To compute similarities between mutations, we used the fitness values where pairs of mutations *g*_*i*_, *g*_*j*_ are measured in matching backgrounds *g*_*k*_ ∈ *N*, allowing us to compute Pearson correlation *ρ*(*g*_*i*_, *g*_*j*_) = *ρ*((*F*(*g*_*i*_, *g*_*k*_) |*g*_*k*_ ∈ *N*), (*F*(*g*_*j*_, *g*_*k*_) |*g*_*k*_ ∈ *N*)). The latter allows us to construct one fully connected graph of mutations where the edges are weighted by the correlation coefficient *A*_*ij*_ = *ρ*(*g*_*i*_, *g*_*j*_). We set a limit of a least 10 the number of matching genetic backgrounds so that the Pearson correlation is more robust. Then, we computed the normalized Laplacian *L*_*n*_ = *I*− *D*^−1*/*2^*AD*^−1*/*2^, where *I* is the identity and *D* is the degree matrix. From *L*_*n*_, we computed the eigenvector corresponding to the smallest non-zero eigenvalue *v*^*F*^, called Fiedler vector. This Fiedler vector encapsulates information about the connectivity of the underlying graph [23]. We initialized the genotype-to-phenotype map *φ* with the values in *v*^*F*^, from which we trained D-LIM at predicting the fitness. This procedure is sensitive to the similarity measure chosen because they capture different relationships (see SI5.1). Therefore, we used the second eigenvalue λ_2_, referred to as the spectral gap, to filter the cases where the spectral initialization is inadequate. For example, when all similarities are *A*_*ij*_ = 1, we have λ_2_ = 1; whereas when all similarities *A*_*ij*_ = 0, we have λ_2_ = 0. We therefore defined the following range λ_2_ > 0.95 or λ_2_ < 0.01 to spectracl initialization, otherwise we use the random Xavier-Glorot method [24].

Correlation between mutations in matching genetic backgrounds. However, because of the nature of Pearson correlation, some landscapes are typically not well processed. For example, the Pearson correlation between two mutations on an additive landscape is always 1, therefore, the similarity matrix is full of ones, which does not inform us about relationships between mutations. In contrast, similarity measured with cosine similarity or Euclidean distance can distinguish mutations in the additive landscape. However, our tests showed that Pearson correlation performs in general better than the other two:

- Cosine similarity is sim 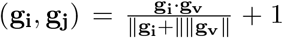, where **g**_**i**_ and **g**_**j**_ are vectors representing fitness values across matching background environments.
- Euclidean similarity is *E*(**g**_**i**_, **g**_**j**_) = exp(−|**g**_**i**_ − **g**_**j**_|).

However, there are cases of symmetric landscapes as shown in Fig SI7, where no similarity measure can differentiate two mutations.

To detect when the similarity measure is inadequate, we use the spectral gap *λ*_2_ (the second eigenvalue) which is a measure of graph connectivity. In our case, it measures how structured is the data (meaning the presence of clusters). An ill similarity matrix where all values tend to 1 has a *λ*_2_ = 1 whereas totally dissimilar mutations with a similarity matrix of zeros yield *λ*_2_ = 0. We therefore defined a range of acceptable *λ*_2_ ∈ [0.05, 0.95].

### 5.2 Models comparison

- **Additive latent model (ALM):** This model used an unconstrained additive latent space, similar to the one proposed in LANTERN. However, here, we used a neural network to predict fitness out of the latent space. Here, we used an 8-dimension latent space, 2 layers of 32-units neural network for fitness prediction. We trained the model for 300 steps, using ADAM and a learning rate of 0.01 and a regularization strength of 0.001. The batch size used here is 64.
- **LANTERN:** This model used an additive latent space, where the importance of each dimension is controlled explicitly. The fitness is predicted using the Gaussian process. Here, we used *K* = 4 latent dimension and a neighborhood number of 200, optimized for 10^3^ steps using ADAM and a learning rate of 0.001.
- **MAVE-NN:** this model introduced a biophysical interpretation using a latent phenotype and a deterministic genotype to phenotype map, combined with a stochastic measurement process. Here, we used an additive genotype-to-phenotype map along with the skewed-t noise model for the measurement process. The model has been trained for 10^3^ steps with a learning rate of 0.001 and batch size of 64.
- **Linear regression (LR):** the linear regression is an additive map we implemented using Pytorch. We optimized the model using Adam for 300 steps with a learning rate of 0.01 and a batch size of 64.

### 5.3 D-LIM hyperparameter and loss function

To train D-LIM, we use the Gaussian negative log likelihood:

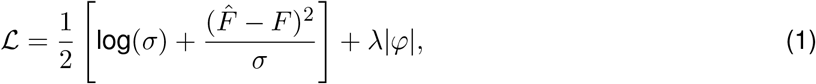

where 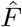 and *σ* are D-LIM predictions respectively for the fitness and its uncertainty, while *λ* = 0.01 regularizes the phenotype values. To choose the hyperparameters for D-LIM, we performed a grid search (for the dataset in [9]), where we varied the number of hidden layers in [1, 2, 3, 4], and the number of neurons per layer in [2, 4, 6, 8, 16, 32]. Based on this search, we selected the architecture with 1 hidden layer of 32 neurons for D-LIM.

We trained D-LIM on a machine equipped with 104 Intel(R) Xeon(R) Gold 6230R CPUs at frequency 2.1HZ, where the learning typically finished in 15.62 seconds for 300 learning steps, see comparisons in Table SI1. We implemented D-LIM using the deep learning backend PyTorch [25], while the analyses were performed using the routines implemented in the package Numpy [26].

## 6 Availability of data and material

Code and data generated as well as fetch from other studies are available in the Github repository https://github.com/LBiophyEvo/D-LIM-model.git. D-LIM is provided in the format of a Python (version 3.10) package. The training time of the model scales linearly with the number of data points, which in the case of [9] is a couple of seconds.

## 8 Supporting information

### 8.1 Cross validation and statistical analyses

All models were evaluated with 30 random fold cross validation where 30% of the original dataset (Env1 in [9]) was left for validation across the whole experiment. The accuracy across folds was quantified with Pearson correlation; whereas, we used the Mean Square Error (MSE) to estimate the gain in predictive power in the context of the extrapolation. Then, to measure the significance of the Pearson correlation sample across random samples and methods, we used the student statistical test (t-test).

To test the generalization of each model, we varied the size of the training set in [99, 152, 232, 354, 540, 824, 999] data points, where we trained 30 model (from random parameters initialization), while the validation set was kept fixed at 30% of the original data.

### 8.2 Kemble *et al* dataset and mechanistic model

The data published in [9] tested all mutation combinations on the promoter of 2 adjacent genes from the linear Escherichia coli l-arabinose pathway. Mutations are restricted to 12 positions in both promoters. There are 72 single mutants and 1296 double mutants, in total 1369 variants. The fitness was measured using barcode tracking in three distinct environments, each defined by specific inducer concentrations: Env1: 20 *ng/ml* aTc and 30 *µM* IPTG; Env2: 5 *ng/ml* aTc and no IPTG; Env3: 200 *ng/ml* aTc and no IPTG). Hence, there are 1369 data samples in each environment.

The mechanistic model proposed in [9] is as follow:

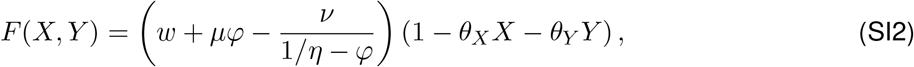

where, 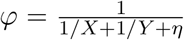 denotes for flux, *η* is the inverse of the maximal flux *φ, θ*_*X*_ and *θ*_*Y*_ represent the cost of increasing cellular enzyme activity, *w* describes the growth rate, *µ* and *ν* are two variables related to downstream enzyme properties.

#### 8.2.1 Extrapolation by using latent variables

We tested here the extrapolation strategy using a non-monotonous fitness function. First, we sampled from the uniform distribution: *ϕ*_1_, *ϕ*_2_ ∼ *U*_[0,5]_, *i* ∈ *{*1, 2 · · ·, 36*}*, representing the true phenotype values. To validate the trained models, we extracted the data points whose phenotype values lay in *ϕ*_1_ ∈ [1.5, 2.7] and *ϕ*_2_ ∈ [1.5, 2.7]. The other data points were used for training, as shown in Fig SI3.

**Table SI1:** CPU time for different algorithms in fitness prediction in Env1 of [9]. Each model was run five times and the results were average (in seconds)

| Models | CPU time (s) |
| --- | --- |
| D-LIM | $15.62 \pm 0.10$ |
| LR | $4.04 \pm 0.01$ |
| ALM | $14.40 \pm 1.93$ |
| Lantern | **$133.90 \pm 135.91$** |
| MAVE-NN | $21.99 \pm 2.63$ |

**Fig. SI1:**
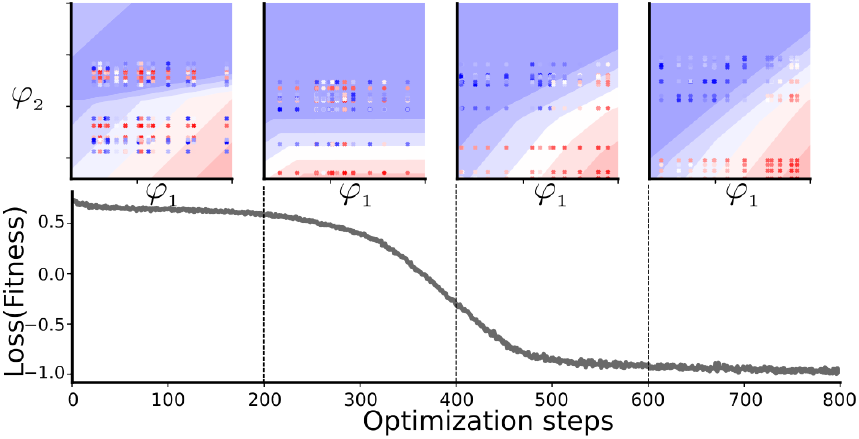
Simultaneous fitting of Fitness and inference of phenotypes. For the mechanistic model [9], we created artificial mutations that were randomly initialized in the inferred phenotype space of D-LIM (first of the four top sub-panels). The real fitness values of each mutant (represented by the color of the dots) did not align with the landscape predicted by D-LIM (the blue-red gradient color); because neither of the parameters of the fitness predictor, nor the inferred phenotype were yet optimized. We monitored the evolution of fitness prediction quality (Loss(Fitness)) and the inferred phenotypes across three-time points, specifically after 200 steps of ADAM optimization. During this optimization process, both the parameters of the fitness predictor and the coordinates in the latent space were adjusted and refined.

**Fig. SI2:**
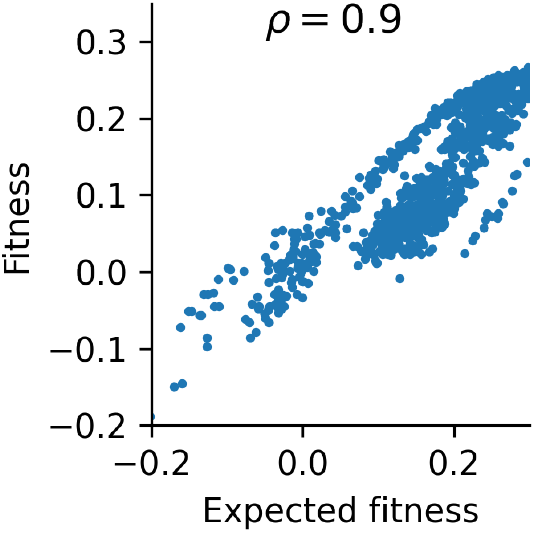
The fitness measured in [9] is largely additive. We compared the double mutant fitness measured F(A, B) with the sum of single mutant fitness F(A)+F(B).

**Fig. SI3:**
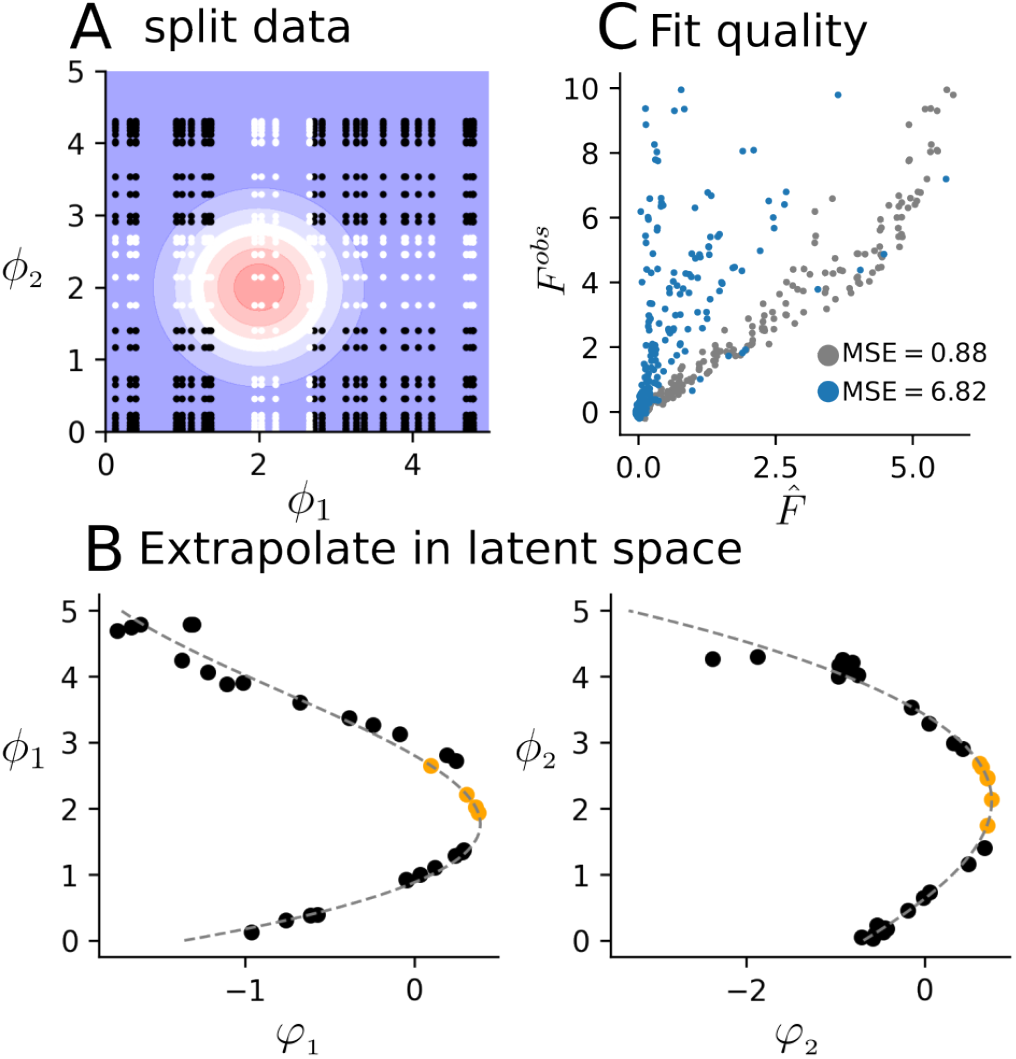
Extrapolation of a non-monotonic fitness landscape. A) Gaussian-type fitness landscape centered at *ϕ*_1_ = 2 and *ϕ*_2_ = 2. Black points are the training points whereas the white ones are the validation points. B) Extrapolation using new *ϕ*_1_ (left), *ϕ*_2_ (right) phenotypic measurements. Black points are (*ϕ*_1_, *φ*_1_) pairs used to parametrize the polynomial fit in a dotted grey line. Orange points are the converted *ϕ*_1_ and *ϕ*_2_ only seen in the validation, which are converted into latent variables using the polynomial fit. C) Fit quality. After converting the phenotypic values only seen in the validation to latent parameters: blue points are the conversion whereas grey ones are after.

**Fig. SI4:**
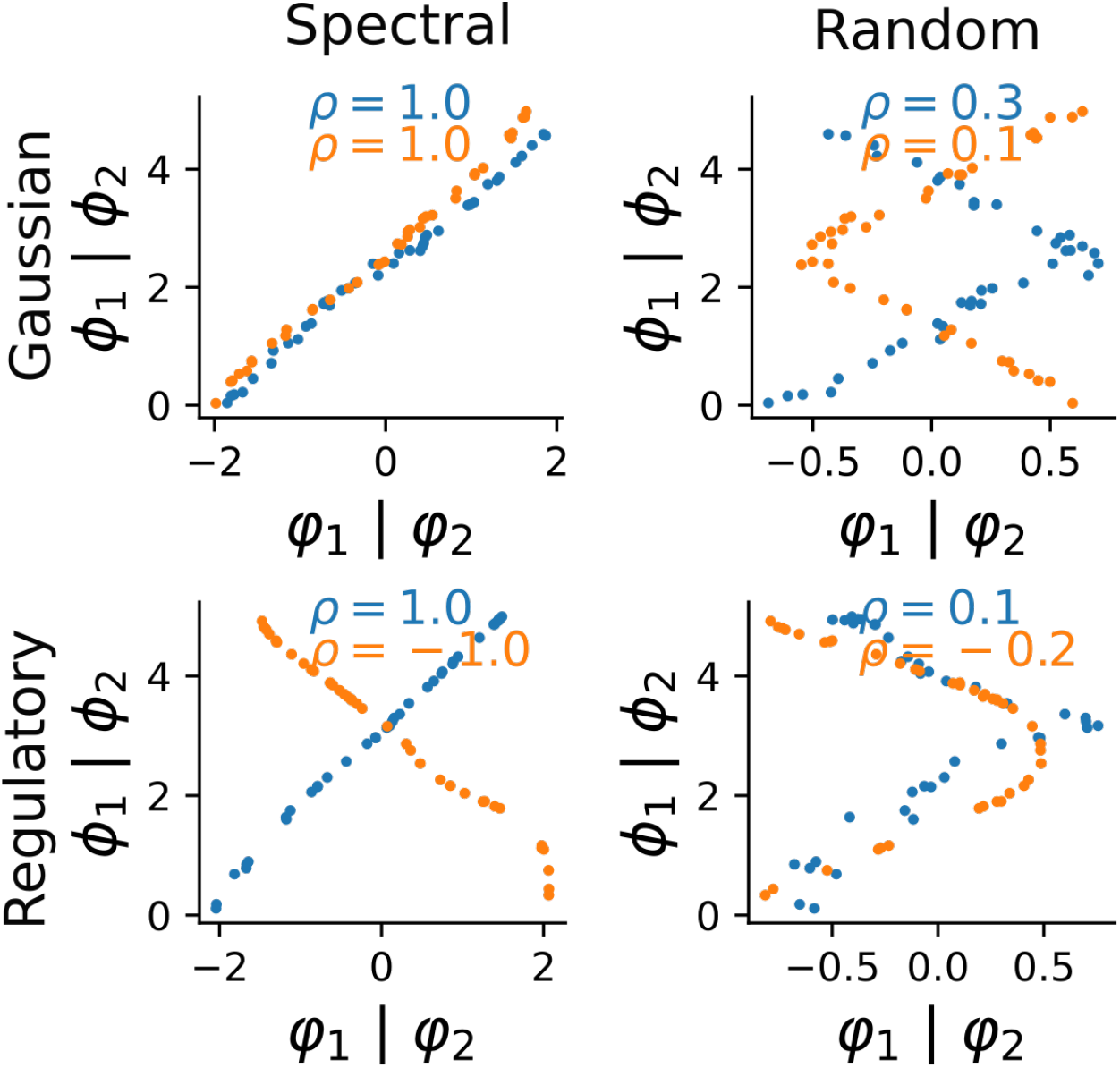
Phenotype inferred after spectral graph initialization for the Gaussian and the regulatory cascade models.

**Fig. SI5:**
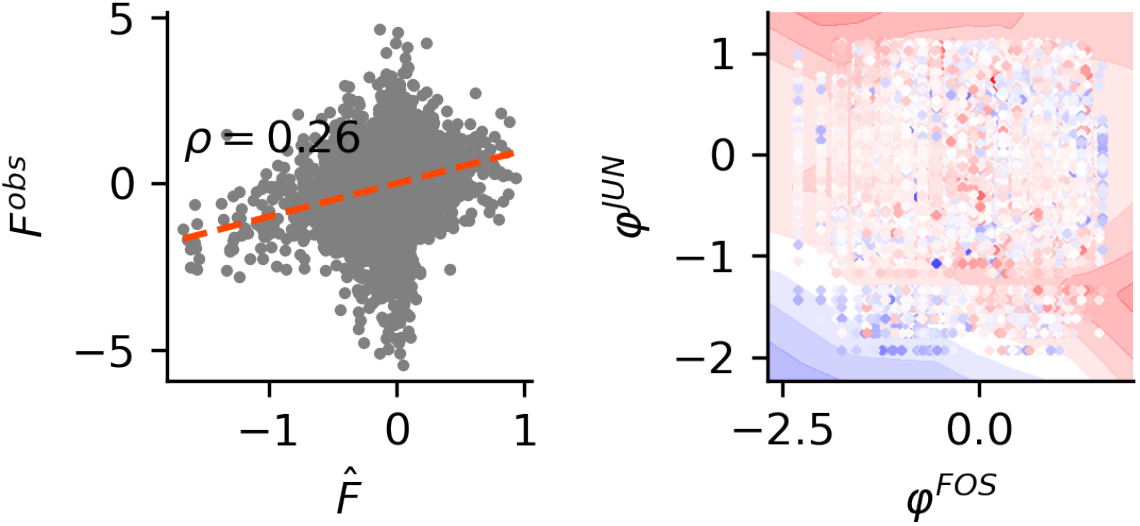
Protein-protein epistasis modeling. The left panel shows the performance of D-LIM for epistasis prediction. The right panel shows the inferred landscape with respect to the D-LIM phenotypes *φ*^*JUN*^, *φ*^*FOS*^.

**Fig. SI6:**
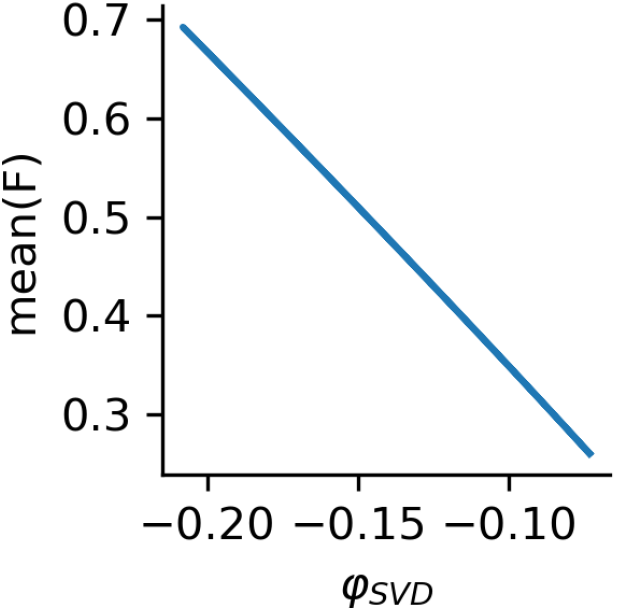
First eigenvector *φ*_*SV D*_ of the SVD applied to the fitness data. Using simulated data from the Fisher model, we compute the *φ*_*SV D*_ and compared it to the mean fitness (mean(F)) across mutations.

**Fig. SI7:**
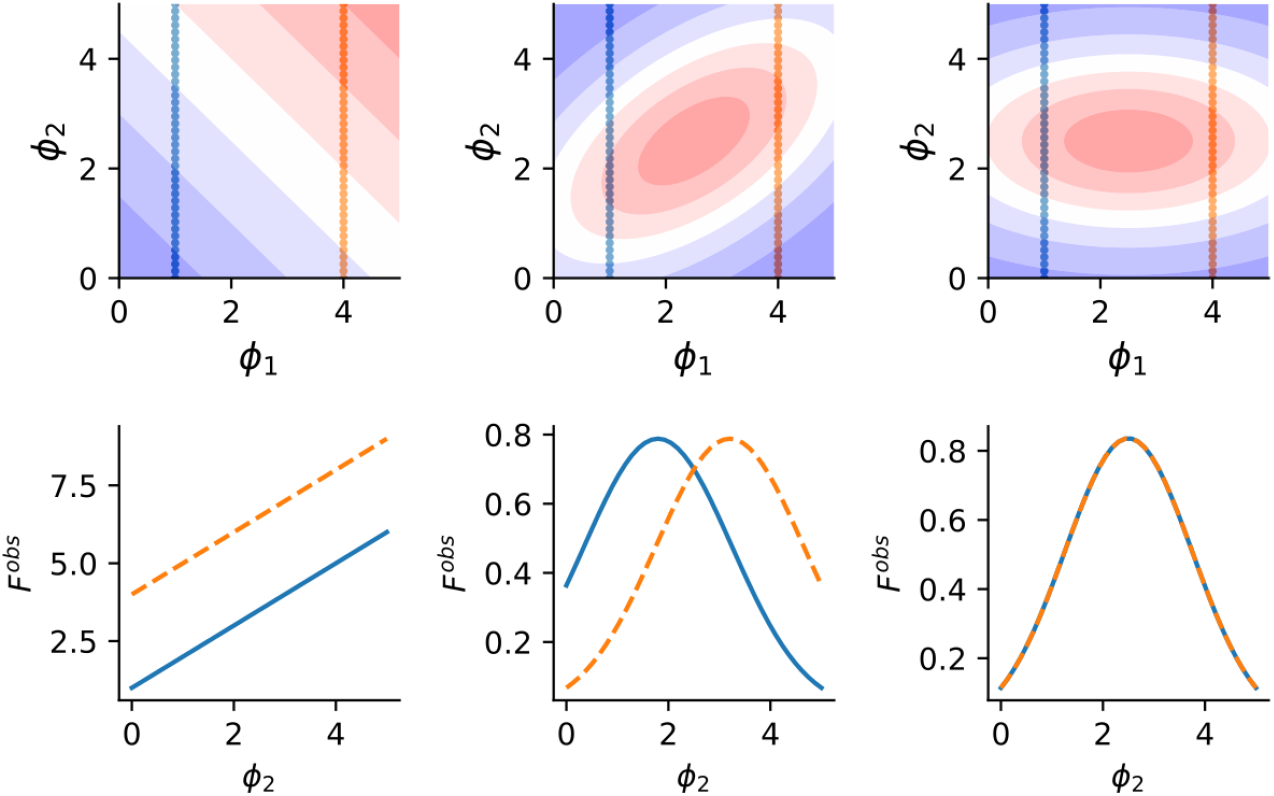
Similarity measures across landscapes. (left) The contour shows the additive model landscape (in the top) where two mutations in *ϕ*_1_ are delineated in blue and orange. Just below, we show the fitness profile obtained across genetic backgrounds. In the center and right, we show the same configuration but on one rotated Fisher model and a non-rotated one.

